# Relationship between birth weight and chronic kidney disease: an integrative analysis of observational studies and causal inference through genetic approaches

**DOI:** 10.1101/729715

**Authors:** Xinghao Yu, Zhongshang Yuan, Haimiao Chen, Jiaji Yang, Yixin Gao, Fengjun Guan, Ping Zeng, Shuiping Huang

## Abstract

**Objective:** Although many observational studies have shown that there was an inverse association between birth weight and chronic kidney disease (CKD) in adults, whether such association is causal remains largely unclear.

**Methods:** We first conducted a systematic review and meta-analysis to investigate the association between birth weight and CKD. Then using a set of valid instrumental variables for birth weight, we performed a two-sample Mendelian randomization (MR) to evaluate its causal effect on CKD based on summary association statistics available from large scale genome-wide association study (GWAS) (up to 143,677 individuals for birth weight and 118,147 individuals for CKD). We further validated the MR results with extensive sensitive analyses.

**Results:** The results of meta-analysis showed that individuals with low birth weight have about 76% (95% CI 36∼126%) higher risk of CKD in late life compared with those with normal birth weight. Depending on 26 instrumental variables, the inverse variance weighted MR showed that the odds ratio per one SD increase of birth weight on CKD was estimated to be 0.91 (95% CI 0.72∼1.14, *p*=0.396). The similar null association between birth weight and CKD is also observed using the weighted median method and maximum likelihood method as well as the Egger regression. Such non-significant association is robust against potential instrumental outliers and pleiotropic effects.

**Conclusion:** Our study identifies an inverse association between birth weight and adult CKD in observational studies, while it is not supportive of the causal role of birth weight on CKD based on our MR analysis.

## 1. Introduction

Chronic kidney disease (CKD) is a common complex disease which influences both children and adult populations (Levey et al., 2015; Webster et al., 2017). At the initial disease stage, CKD is asymptomatic and may be ignored by sufferers. It is common that the diagnosis of CKD is made when disease symptoms already become severe. Moreover, a series of severe complications (e.g. renal failure, hypertension, cancer, infection and coronary heart disease) can occur with the decreased renal function of CKD patients (Di Lullo et al., 2015; Webster et al., 2017). Additionally, due to the decreased GFR during the disease progression as well as possible complications, the life quality of CKD patients is significantly lower than that of the general population.

It is estimated that the global prevalence of CKD ranges between 11% and 13%, and that CKD can account for 1.5% death worldwide, making it among the leading death risk and a global public health issue (Hill et al., 2016). World Health Organization (WHO) predicts that the deaths attributable to kidney-related diseases will increase by 31% (from ∼871,000 in 2015 to ∼1,143,000 in 2030) in the next decade due to the growing disease rate and aging population (Organization, 2018). Therefore, identifying the risk factors of CKD can promote our understanding of the pathogenesis of this disease, having the potential to ultimately lead to better prevention and treatment for CKD, and is also important in terms of the public health perspective (Luyckx and Brenner, 2015; Luyckx et al., 2017; Webster et al., 2017).

CKD has complicated etiologies and both genetic and non-genetic (e.g. lifestyle and environmental) risk factors to play an important role in the development of CKD (Jha et al., 2013; Webster et al., 2017; Iwagami et al., 2018; Wang et al., 2018). Some non-genetic risk factors (e.g. diabetes, hypertension, dyslipidemia and glomerulonephritis) were previously discovered in observational studies. Additionally, multiple genes (e.g. *NAT8, SLC7A9, UMOD, SHROOM3, GATM* and *MYH9*) were also identified to be associated with CKD and kidney-related traits (Chambers et al., 2010; Köttgen et al., 2010; Pattaro et al., 2016). More recently, several epidemiological studies have shown that CKD may originate from the life of the fetus —— a generalized hypothesis referred to as the fetal origins hypothesis first proposed by the British epidemiologist David Barker in 1990 (thus also known as Barker hypothesis) (Barker, 1990; Luyckx and Brenner, 2015). The fetal origins hypothesis supposes that the risk for chronic non-communicable diseases (e.g. CKD and cardiovascular diseases) in later life can be partly attributed to the altered developmental programming and the long-term adverse adaptations to early undernutrition, both of which can lead to the structural and functional changes in multiple developing tissues and organs (Zeng et al., 2019b; Zeng and Zhou, 2019a). In the literature, birth weight is a widely used measurement for intrauterine environment; and low birth weight often serves as an indicator of impaired renal development in utero when investigating the influence of early growth on kidney-related outcomes. Although most previous studies (White et al., 2009; Luyckx and Brenner, 2015; Das et al., 2016), along with some animal experimental models (Barnett et al., 2017), illustrated that low birth weight was associated with an increased risk of CKD; owing to the heterogeneity in disease onset age, geographic diversity and ethnic differences, a few of other studies did not support the existence of the inverse relationship between birth weight and CKD (Fagerudd et al., 2006; Haysom et al., 2007), and sometimes even reported contradictory findings (Vasarhelyi et al., 2000). For example, no early glomerular and tubular damage was observed in young men with low birth weight compared with those with normal birth weight (Vasarhelyi et al., 2000).

The inconsistent observations regarding the relationship between birth weight and CKD may be also partly due to uncontrolled/unknown confounders which are commonly encountered in observational studies. Indeed, there are studies which suggested that the impaired kidney function in adulthood may be a consequence of high blood pressure (Vasarhelyi et al., 2000). Thus, it remains a concern when interpreting the observed relationship between birth weight and CKD as a causal association. A cohort longitudinal study may alleviate such concern and offer an important insight into the causal interpretation. However, longitudinal studies require large scale subjects and need very long-term follow-ups before CKD clinical presentation. Traditionally, randomized controlled trials (RCT) studies are the gold standard for inferring the causal effect of exposure on outcome. However, determining the causal relationship between birth weight and CKD by RCT is infeasible. It seems that the validation of the fetal origins hypothesis for CKD is extremely difficult in a traditional manner.

In observational studies Mendelian randomization (MR) can help clarify the causal relationship between an exposure of interest and an outcome, and provide an efficient way for causal inference. Briefly, MR is a special instrumental variable method that employs genetic variants (e.g. single nucleotide polymorphisms (SNPs)) as instruments for an exposure (i.e. birth weight) and evaluates its causal effect on the outcome (i.e. CKD). In the past ten years the great success of genome-wide association studies (GWASs) makes it feasible to select suitable SNPs as effective instruments for causal inference in MR. In fact, MR has recently become a considerably popular approach of inferring causal relationship in observational research (Mokry et al., 2015; Zeng et al., 2019a; Zeng and Zhou, 2019b). Indeed, birth weight has been confirmed to be causally associated with many adult diseases (e.g. cardiovascular disease (Au Yeung et al., 2016; Zanetti et al., 2018) and type 2 diabetes (Wang et al., 2016)) through MR studies.

Motivated by those previous observations above, our main goal in this study was two aspects. First, to illuminate whether there exists an association between birth weight and CKD, we employed the systematic review and meta-analysis to provide a pooled conclusion. The result showed that birth weight is inversely associated with CKD, confirming the finding in other studies (Lackland et al., 2000; Fan et al., 2006; Al Salmi et al., 2008; Oster et al., 2013; Hirano et al., 2016). Furthermore, to determine whether this observed negative association is causal, we performed a largest and most comprehensive MR analysis based on summary statistic data available from large-scale GWASs with approximately 143,000 individuals for birth weight and ∼118,000 individuals for CKD.

## Materials and Methods

### Systematic reviews and meta-analysis

#### Data sources and search strategies for previous studies

Following the guideline of preferred reporting items for systematic reviews and meta-analyses (PRISMA) (Moher et al., 2009), we performed a literature search mainly on PubMed from January 1998 to April 2019 for articles on the relationship between birth weight (and related factors including premature birth and fetal development) and CKD. We made no restriction on study designs and considered both cohort and population-based studies; but we limited articles in English. Originally, a total of 2,072 articles (2,067 articles by searching and additional 5 articles by references scanning) were obtained (Fig. S1). The following exclusion criteria for articles filtering were employed: (1) the title and abstract did not contain any data on birth weight and/or CKD; (2) insufficient results were available on birth weight and CKD; (3) duplicated studies; (4) articles were a review, letter-to-editor, response or commentary article; and (5) articles were about clinical drug trials for CKD; (6) articles were about CKD in childhood. Based on those criteria, 20 studies were left in our final meta-analysis.

#### Data extraction and Statistical analysis in meta-analysis

For each article that was incorporated into our meta-analysis, two investigators (XH and PZ) independently carried out data extraction and quality assessment. From each article we extracted the information about study setting and design, population and sample size for case and control, effect size (e.g. odds ratio (OR), relative ratio (RR) or hazard ratio (HR)) as well as covariates that were adjusted for in the original analysis. The effect size heterogeneity among studies was tested by the Cochran’s Q statistic (Thompson and Sharp, 1999). We estimated the combined effect of birth weight on CKD using a weighted meta-analysis method and evaluated the published bias by the Egger method and funnel plot (Egger et al., 1997). We also performed a leave-one-out (LOO) analysis to assess the influence of a single study.

### MR analysis

#### GWAS data sources for birth weight and CKD

In our meta-analysis above we found that there exists a robust inverse association between birth weight and CKD (see below for more details). To examine whether this identified association is causal, we further performed a MR analysis based on large scale GWAS genetic data of birth weight and CKD. To achieve this, we first yielded the genetic data of birth weight from the Early Growth Genetics (EGG) consortium (http://egg-consortium.org) (Horikoshi et al., 2016). In this study, birth weight was measured as a continuous variable and an additive linear regression was adopted for each SNP to detect its association with birth weight while controlling for available covariates (e.g. gestational age). After quality control of SNP genotypes and individuals, it contained summary association statistics (e.g. effect allele, marginal effect size, standard error, p value and sample size) for 16,245,523 genotyped and imputed SNPs on 143,677 individuals of European ancestry.

We next obtained the summary association statistics (e.g. marginal effect size, standard error and p value) of CKD from the CKDGen consortium (http://ckdgen.imbi.uni-freiburg.de/) (Pattaro et al., 2016). After quality control a total of 118,147 European individuals (12,385 cases and 105,762 controls) and 2,191,883 SNPs were available for this CKD GWAS. Besides CKD, we also attempted to explore the causal relationship between birth weight and other kidney-related phenotypes which included eGFR based on serum creatinine (eGFRcrea) and cystatin C (eGFRcys) (Pattaro et al., 2016), annual decline of eGFR (eGFR change) and rapid eGFR decline (Rapid Decline) (Gorski et al., 2015), urinary albumin-to-creatinine ratio (UACR) and microalbuminuria (MA) (Teumer et al., 2016). The used GWAS genetic data sets in our MR study are summarized in Table S1. Since participants had given informed consent for data sharing as described in each of the original GWASs and only summary association results were employed; therefore, ethical review was not needed for our study.

#### Estimation of causal effect of birth weight on CKD and sensitivity analyses

We then employed MR approaches to determine the causal relationship between birth weight and CKD. First, to ensure the validity of MR we carefully selected a set of independent index associated (*p*<5.00E-8) SNPs that can serve as valid instrumental variables for birth weight. The summary information of those index SNPs for birth weight and CKD are shown in Table S2. Next, to quantitatively check whether the selected instruments for birth weight are strong, we calculated the proportion of phenotypic variance of birth weight explained by each instrument and computed the *F* statistic as an empirical indicator of strong/weak instrument (Noyce et al., 2017). We then performed the two-sample inverse-variance weighted (IVW) MR methods (Burgess et al., 2017) to estimate the causal effect of birth weight on CKD in terms of per standard deviation (SD) change in birth weight, where the SD of birth weight was estimated to be about 488 grams (Horikoshi et al., 2016). Before the causal effect estimation, to further ensure the validity of MR, we inspected the pleiotropic associations by removing instruments that may be potentially related to CKD with an adjusted p value less than 0.05 after Bonferroni correction. In our analysis no instruments were excluded by this conservative manner. To examine the robustness of results in the MR analysis, we carried out several sensitivity analyses: (1) weighted median-based method (Bowden et al., 2016) and maximum likelihood method (Burgess et al., 2013); (2) LOO analysis (Noyce et al., 2017) and MR-PRESSO test (Verbanck et al., 2018) to validate instrumental outliers that can substantially impact the causal effect estimate; (3) MR-Egger regression to detect directional pleiotropic effects of instrument variables (Burgess and Thompson, 2017).

## Result

### Combined effect of birth weight on CKD in systematic review and meta-analysis

A total of 20 studies satisfied the inclusion criteria and were finally incorporated into our systematic review and meta-analysis (Fig. S1). Most of the included studies were carried out on European individuals. The extracted information of those studies is shown in Table 1. All the studies reported the risk of CKD for low birth weight compared with normal birth weight, and nine additionally reported the risk of CKD for high birth weight compared with normal birth weight. Note that the definition of low/high birth weight was slightly different across studies (see Table 1 for more details). Among those, 16 studies showed that low birth weight can increase the risk of CKD in later life. Additionally, five out of nine studies demonstrated that high birth weight can also raise the risk of CKD. Those results suggested that there may exist a U-shaped relationship between birth weight and CKD. We thus performed meta-analysis for the association between low or high birth weight with CKD separately (Fig. 1).

**Table 1.**
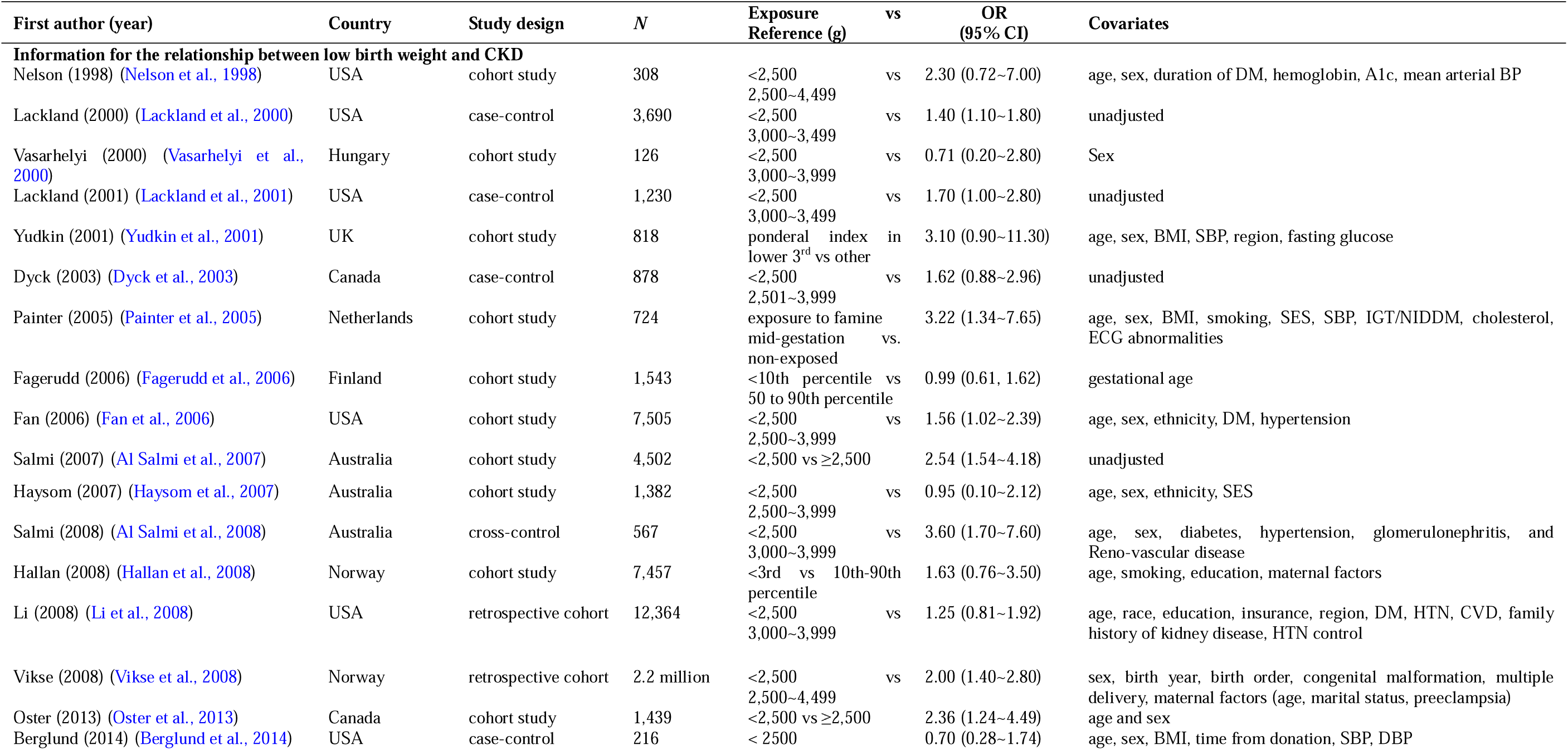
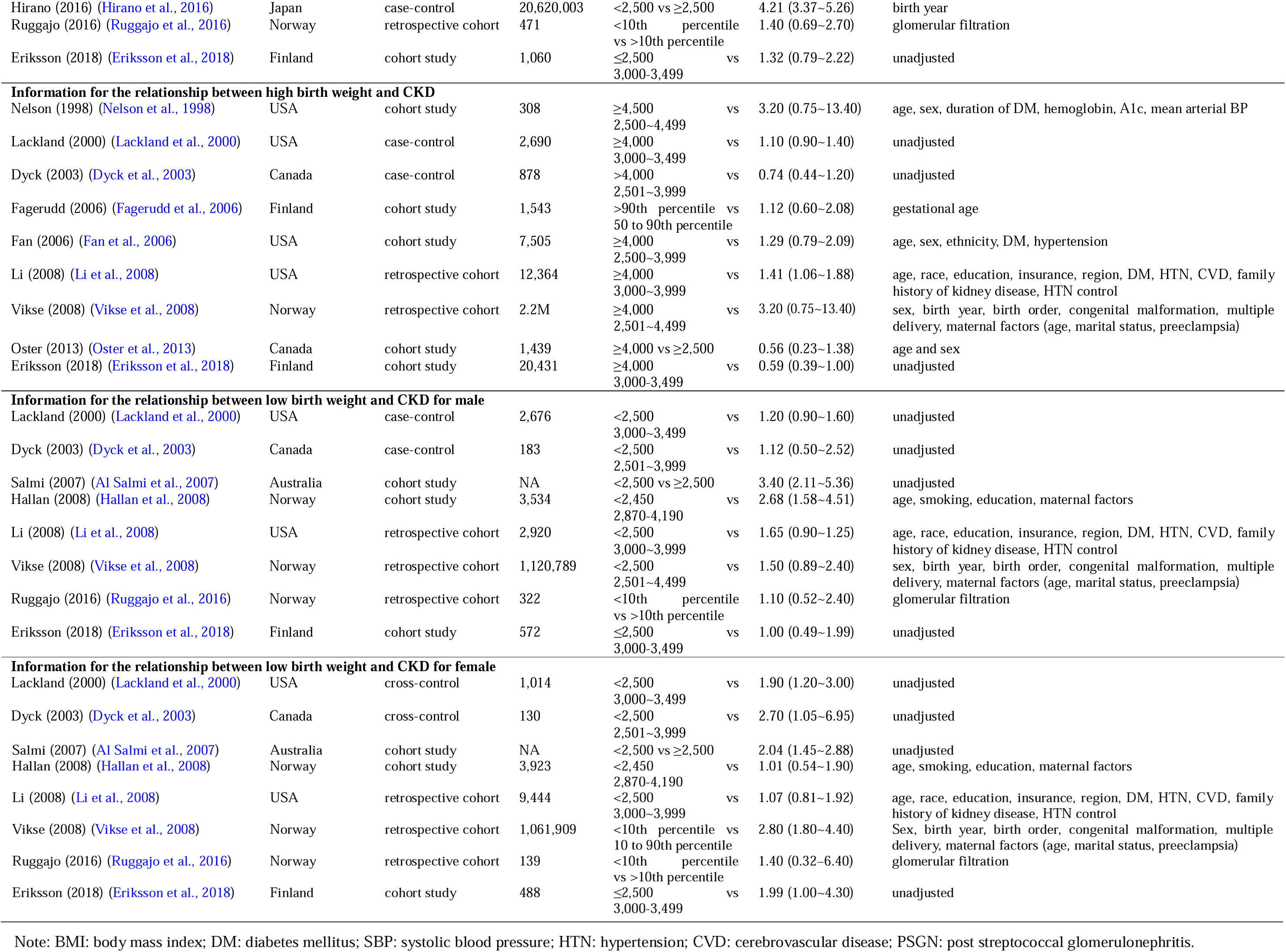
Summary information of 20 studies included in the meta-analysis for investigating the relationship between birth weight and chronic kidney disease

**Fig. 1.**
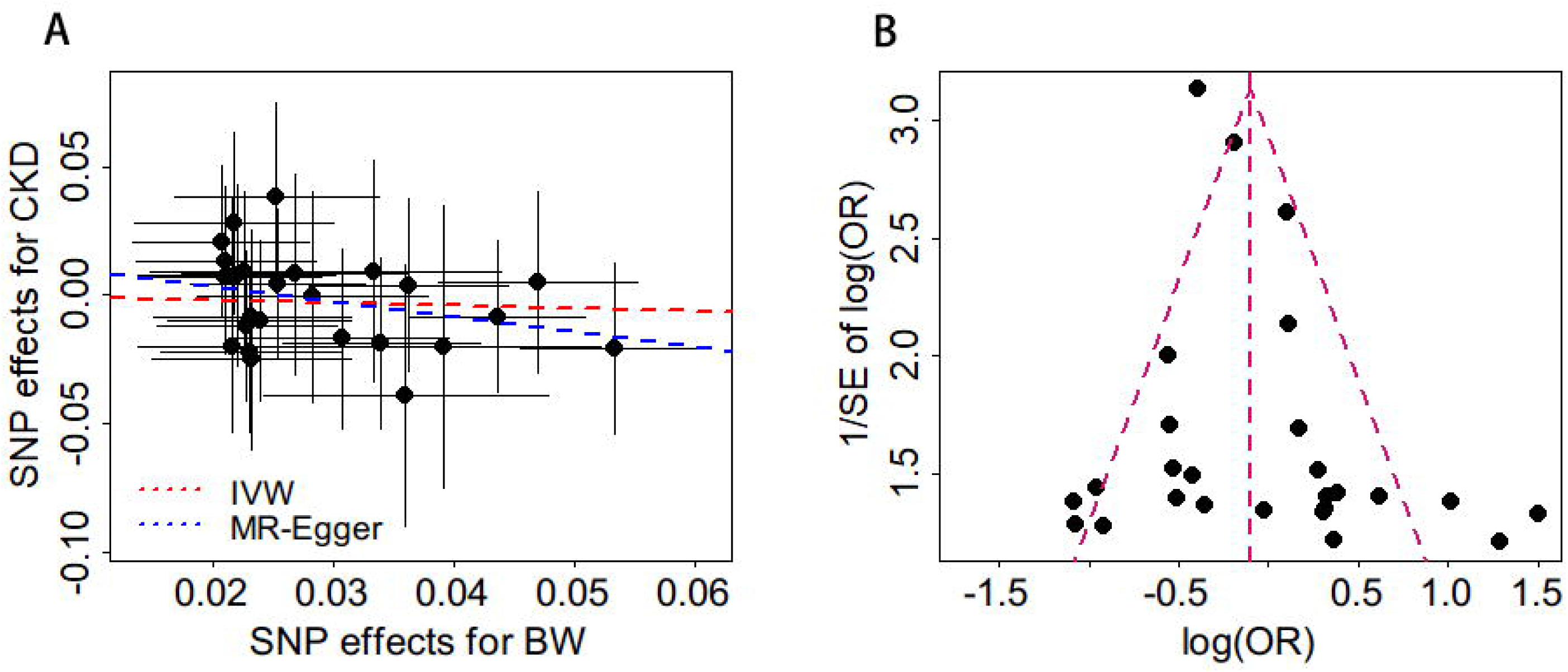
Combined effect of birth weight on CKD in the meta-analysis based on observational studies. (A) Combined effect for individuals with low birth weight compared with those with normal birth weight based on twenty studies; (B) Combined effect for individuals with high birth weight compared with those with normal birth weight based on nine studies.

Owing to the presence of heterogeneous effect size of birth weight on CKD in those studies (the p values of the Q statistic are less than 0.05 for both low and high birth weight; Fig. 1), the results of the random-effects meta-analysis are displayed here. Specifically, we found that the risk of CKD for adult individuals with low birth weight is 76% (OR=1.76, 95% CI 1.37∼2.26, *p*=1.27E-5) higher compared with those with normal birth weight (Fig. 1A), implying that lower birth weight leads to more vulnerable to CKD. This inverse relationship also holds in the sub-group meta-analyses in terms of gender or the type of study design (Fig. S2-S3). However, no significant association is observed between high birth weight and CKD (OR=1.05, 95% CI 0.81∼1.37, *p*=0.713; Fig. 1B). These results are robust according to the LOO analyses which show that no single study can substantially dominate the final combined estimates (Table S3-S4). Additionally, the Egger test (*p*=0.170 for low birth weight and *p*=0.982 for high birth weight), together with the funnel plot (Fig. S2), demonstrates that the publication bias is less likely to influence the combined estimates in our meta-analysis. In summary, based on the results of meta-analysis above, we can conclude that an inverse association exists between birth weight and CKD, but no evidence is present for the observed U-shaped relationship.

### Estimated causal effect of birth weight on CKD

In our MR analysis, a total of 26 independent index SNPs served as instrument variables for birth weight. They jointly explain a total of 0.91% of phenotypic variance for birth weight. The *F* statistics of those instruments range from 27.6 to 175.6 (with an average of 49.26), indicating that the weak instrument bias does not likely occur in our analysis. Little evidence of causal effect heterogeneity across instruments is observed (Q=23.08 and *p*=0.573); therefore, we employed the fixed-effects IVW MR method to estimate the causal effect and found that there exists a negative but non-significant casual association between birth weight and CKD. More specifically, the OR per one SD increase of birth weight on CKD is 0.91 (95% CI 0.72∼1.14, *p*=0.396), consistent with those produced by the weighted median method (OR=0.86, 95% CI 0.62∼1.18, *p*=0.346) and by the maximum likelihood approach (OR=0.91, 95% CI 0.72∼1.14, *p*=0.414). The similarly null causal association was also observed if we employ other sets of instrumental variables for birth weight (Supplementary File). In addition, we also did not discover a significant casual association between birth weight and other kidney-related traits (Fig. S5).

We next examined whether there are potential instrument outliers and whether these outliers have a substantial influence on the estimate of causal effect. To do so, we created a scatter plot by drawing the effect sizes of SNPs of birth weight against those SNPs of CKD for all the used instruments (Fig. 2A). It is shown that no instrumental variables can be considered potential outliers. The result of MR-PRESSO also displays that there do not exist instrument outliers at the significance level of 0.05. Consistently, in terms of the result of the LOO analysis, no single instrument can have a substantial influence on the estimation of causal effect (Table S5). The OR per one SD increase of birth weight on CKD is estimated to be 0.55 (95% CI 0.26∼1.17, *p*=0.120) using the MR-Egger regression. Furthermore, the MR-Egger regression removes the possibility of pleiotropic effects of instrument variables (the intercept=0.015, 95% CI -0.007∼0.037, *p*=0.174). The funnel plot also presents a symmetric pattern around the causal effect point estimate (Fig. 2B), further indicating the absence of horizontal pleiotropy. Overall, the MR results do not provide statistically significant evidence that supports the direct causal association between birth weight and CKD.

**Fig. 2.**
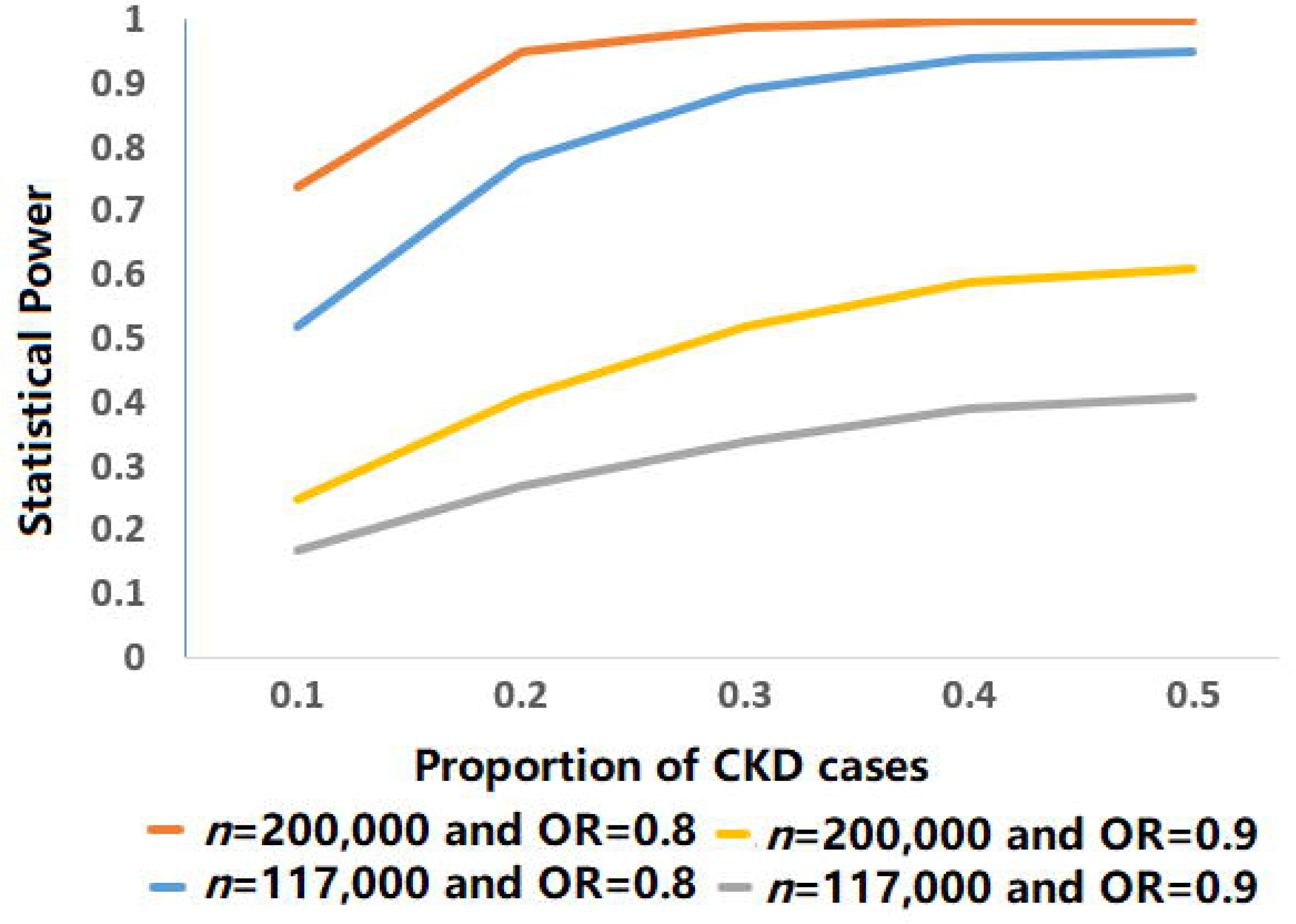
(A) Relationship between the SNP effect size estimates of birth weight (x-axis) and the corresponding effect size estimates of CKD (y-axis). In the plot, the 95% CIs for the effect sizes of instruments on birth weight are shown as horizontal lines, while the 95% CIs for the effect sizes of instruments on CKD are shown as vertical lines. The line in red represents the estimated causal effect of birth weight on CKD obtained using the IVW method while the blue line represents the estimated causal effect produced by the MR-Egger regression. (B) Funnel plot for single causal effect estimate of birth weight on CKD; the horizontal dot line denotes the estimated causal effect with IVW.

## Discussion

To understand the relationship between birth weight and CKD, in the present study we first performed a systematic review and meta-analysis. The results showed that individuals with low birth weight would have a higher risk of CKD in adulthood compared with those with normal weight, in line with previous observation (Poulter et al., 1999; Al Salmi et al., 2007; Khalsa et al., 2016). The mechanism underlying this inverse association between birth weight and CKD is very complicated (Di Lullo et al., 2015; Webster et al., 2017). Possible interpretations include the finding that low birth weight can lead to the reduction of the number of kidney nephrons (Luyckx et al., 2013). For example, it was observed that every 1 kg decrease of birth weight can result in about 250,000 reduction in the number of unilateral nephrons (Hoy et al., 2005). The relatively smaller number of nephrons for individuals with low birth weight implies a higher susceptibility to kidney diseases in later life (Brenner et al., 1988; Luyckx et al., 2017). This finding was also supported by animal models which showed that offspring had decreased kidney nephrons if being exposed to adverse environmental conditions during pregnancy (Bidani et al., 2013; Horowitz et al., 2015). However, our results provided little evidence supporting the existence of association between high birth weight and CKD although previous studies suggested high birth weight can also elevate the risk of CKD.

To investigate whether this observed negative association between birth weight and CKD in our meta-analysis is causal, we further carried out a two-sample MR analysis based on summary statistics publicly available from large scale GWASs. Because MR relies on the Mendel’s second law which means that an allele of a gene can enter a gamete independently of another gene. Therefore, MR is less likely affected by confounding factors compared with observational studies (Burgess et al., 2017). In our MR analysis, to improve the statistical power and meet the model assumptions we used multiple instrument variables which were independent from each other and strongly associated with birth weight. We also tried to avoid the pleiotropic effects of instruments by removing index SNPs that may be potentially related with CKD. Further sensitive analyses (e.g. Egger regression) also excluded the likelihood of pleiotropy that can introduce bias into the causal effect estimation. However, the results of MR did not offer statistically significant evidence supporting the direct causal relationship between birth weight and CKD.

Several explanations exist in this observed association. Especially, after birth the threat to the survival of the nephron still exists. Infants with low birth weight are often accompanied by a decrease in the number of nephrons due to impaired renal function development. The decrease in the number of nephrons may result in glomerular hypertrophy and high filtration rate, which ultimately leads to secondary glomerular sclerosis. As one of the important risk factors for CKD (Coca et al., 2012), acute kidney injury occurs in 18-40% of very low birth weight infants (Koralkar et al., 2011). Additionally, most of infants with very low birth weight receive at least one nephrotoxic drug treatment before discharge, which can potentially affect kidney function (Rhone et al., 2014). Infants with low birth weight or limited intrauterine growth often experience accelerated “catch-up” growth, which is also associated with CKD (Fagerberg et al., 2004).

Nevertheless, we note that the estimated causal effects between birth weight and CKD were consistent in the direction and magnitude through multiple MR methods (e.g. IVW, weighted median method and maximum likelihood estimation). There are several explanations for the failure of detecting a causal association between birth weight and CKD given the observation that low birth weight is robustly related to the increased risk of CKD in our meta-analysis. First, this inverse relationship in observational studies may be driven by shared genetic components between birth weight and CKD. To check this, we applied the linkage disequilibrium score regression (LDSC) (Bulik-Sullivan et al., 2015) to quantify the genetic covariance between birth weight and CKD. LDSC is a novel statistical genetic method for quantifying genetic correlation for two traits based on the genome-wide pleiotropy (note that our MR analysis has removed the influence of pleiotropic effects). With LDSC, we found a pronounced but nonsignificant genetic correlation between birth weight and CKD (*R*_g_=-0.234, se=0.081, *p*=0.771; see Table S6 for more information), suggesting the common polygenic risk shared by low birth weight and CKD. More specifically, this means that some SNPs that are associated with low birth weight also related to the risk of CKD. Second, the failure of detecting non-zero causal effect of birth weight on CKD may be partly due to a lack of adequate statistical power. To examine this, we performed the statistical power calculation to discover an OR of 0.80 or 0.90 in the risk of CKD per unit change of birth weight following the approach shown in (Brion et al., 2013). Note that, these assumed ORs approximately equal to the estimated effect size of birth weight on CKD in our study. The results imply that we have a small to moderate power to detect the causal association between birth weight and CKD due to the small number of CKD cases (Fig. 3). For example, with the current sample size of CKD in our study (i.e. assume the sample size of adult CKD is 117,000 and the proportion of cases is 10.6%), the estimated statistical power is 17% or 25% to detect an OR of 0.80 or 0.90, respectively. Third, we cannot rule out the possibility that there exist some unknown pathways which mediate the influence of birth weight on CKD. Note that the existence of mediation effect (or indirect effect) of birth weight does not violate the model assumptions of MR. For example, it is well-established that low birth weight can increase the risk of coronary heart disease, diabetes and hypertension; the latter two are the major causes of CKD (Wingen et al., 1997; Jafar et al., 2003; Ardissino et al., 2004; Targher et al., 2008; Jha et al., 2013), implying that birth weight can have an impact on CKD by the metabolic or cardiovascular pathway.

**Fig. 3.**
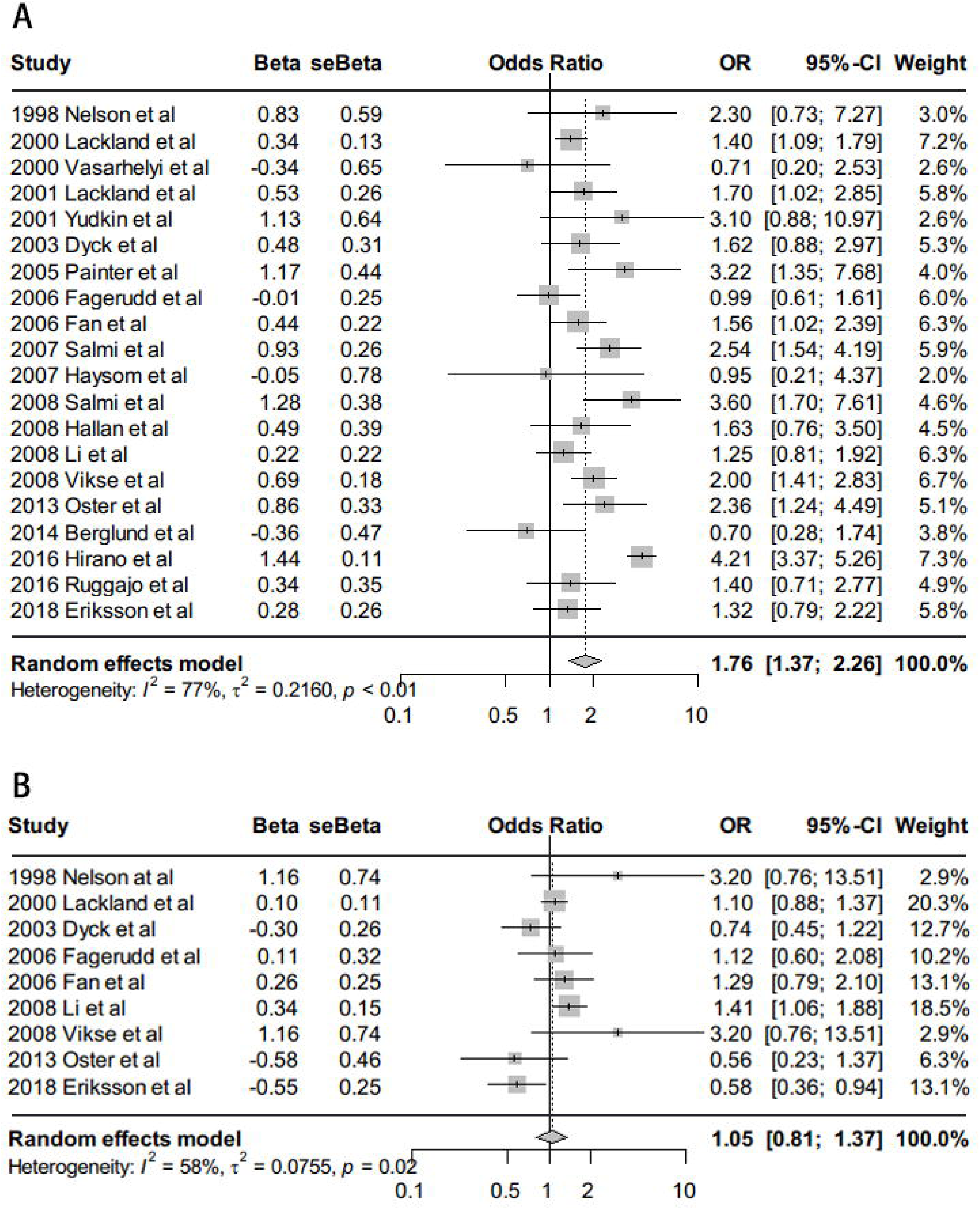
Statistical power estimated with the analytic method shown in (Brion et al., 2013). In the estimation, the total phenotypic variance explained by instrumental variables was set to be 0.91%, the significance level α was set to be 0.05, the proportion of CKD cases was set to be from 0.1 to 0.5. Two situations of sample size (i.e. 117,000 and 200,000) were considered. For each situation, the OR was assumed to be 0.80 or 0.90, respectively.

### Limitation

Finally, some limitations of this study should be considered. First, both birth weight (and all corresponding antecedents and early risk factors) and CKD are heterogeneous phenotypes; for example, among adult CKDs, polycystic kidney disease is currently known as a hereditary kidney disorder and is one of the most common autosomal dominant diseases (Gabow, 1993; Chapman et al., 2015). Diabetic nephropathy is caused by lifestyle and genetic factors and Hypertensive kidney disease is more caused by environmental factors(Go et al., 2004; Vivante and Hildebrandt, 2016). Therefore, when combining these heterogeneous CKDs together in analysis, a large degree of deviation may be introduced in our analysis. Second, as mentioned above, we have only a limited statistical power in our MR analysis due to the small sample size of cases in the CKD GWAS. Third, like many previous MR studies we hypothesized that there is a linear relationship between birth weight and CKD in our analysis. Linearity may be unreasonable in practice since previous epidemiological studies have found that high birth weight also increases the risk of CKD, implying a U-type relationship between birth weight and CKD. Therefore, we cannot completely remove the nonlinear influence of birth weight on CKD. Fourth, our MR relies on summary statistics rather than individual-level data sets, thus we cannot analyze the relationship between very low/high birth weight and CKD due to lack of relevant data information, and we are also unable to conduct stratified analyses (e.g. in terms of gender; see Table 1) in our MR study.

In conclusion, our study identifies an inverse association between birth weight and CKD in observational studies, while it is not supportive of the causal role of birth weight on the risk of CKD based on our MR analysis.

## Acknowledgements

We thank all the EGG and CKDGen consortium studies for making the summary data publicly available and we are grateful of all the investigators and participants contributed to those studies. The data analyses in the present study were supported by the high-performance computing at Xuzhou Medical University.

## Author Contributions

PZ and SH conceived the idea for the study; PZ and XY obtained the data; PZ and XY cleared up the datasets; PZ, XY and ZY mainly performed the data analyses; HC, YG and JY helped clear and analyze the data; PZ, XY, ZY and FG interpreted the results of the data analyses; PZ and XY wrote the manuscript, and other authors approved the manuscript and provided relevant suggestions.

## Disclosure

The authors declare that the research was conducted in the absence of any commercial or financial relationships that could be construed as a potential conflict of interest.

## Funding

This study was supported by the National Natural Science Foundation of Jiangsu (BK20181472), Youth Foundation of Humanity and Social Science funded by Ministry of Education of China (18YJC910002), the China Postdoctoral Science Foundation (2018M630607 and 2019T120465), Jiangsu QingLan Research Project for Outstanding Young Teachers and Six Talent Peaks Project of Jiangsu Province of China (WSN-087), the Postdoctoral Science Foundation of Xuzhou Medical University, the National Natural Science Foundation of China (81402765), the Statistical Science Research Project from National Bureau of Statistics of China (2014LY112), the Postgraduate Research & Practice Innovation Program of Jiangsu Province (KYCX19_2250), and the Priority Academic Program Development of Jiangsu Higher Education Institutions (PAPD) for Xuzhou Medical University.

## Supplementary material

Supplementary File.

